# SCONCE: A method for profiling Copy Number Alterations in Cancer Evolution using Single Cell Whole Genome Sequencing

**DOI:** 10.1101/2021.09.23.461581

**Authors:** Sandra Hui, Rasmus Nielsen

## Abstract

Copy number alterations are a significant driver in cancer growth and development, but remain poorly characterized on the single cell level. Although genome evolution in cancer cells is Markovian through evolutionary time, copy number alterations are not Markovian along the genome. However, existing methods call copy number profiles with Hidden Markov Models or change point detection algorithms based on changes in observed read depth, corrected by genome content, and do not account for the stochastic evolutionary process. We present a theoretical framework to use tumor evolutionary history to accurately call copy number alterations in a principled manner. In order to model the tumor evolutionary process and account for technical noise from low coverage single cell whole genome sequencing data, we developed SCONCE, a method based on a Hidden Markov Model to analyze read depth data from tumor cells using matched normal cells as negative controls. Using a combination of public datasets and simulations, we show SCONCE accurately decodes copy number profiles, with broader implications for understanding tumor evolution. SCONCE is implemented in C++11 and is freely available from https://github.com/NielsenBerkeleyLab/sconce.

## 1. Introduction

In cancerous cells, somatic driver and passenger single nucleotide polymorphisms (SNPs) and copy number alterations (CNAs) accumulate over time. CNAs are extremely common across cancer types (1, 2).

Many large scale cancer studies are done with bulk samples, and many methods and evaluation techniques (3, 4) have been developed to identify copy number alterations in bulk sequencing, especially for low coverage data (5) and tumor heterogeneity deconvolution (6). However, bulk sequencing averages mutations across many cells and loses the granularity and detail single cell sequencing (SCS) can provide. Using single cell sequencing, we can evaluate these mutations on a cell by cell level and treat each cell as an individual in a population. However, the SCS process is technically challenging and produces noisy low coverage data, due to challenges like cell dissociation, small amounts of starting DNA, and non uniform whole genome amplification (7). Although the rapidly increasing availability of single cell RNA sequencing (scRNA-seq) of tumors can yield insights into tumor subpopulations (8) and relevant biological pathways and processes (9, 10), using scRNA-seq for calling CNAs is limited to areas of the genome that are expressed at the time of sequencing and does not directly measure genomic copy number. However, single cell whole genome DNA sequencing data promises to circumvent these problems, despite the inherent noisiness of the data.

The main components of CNA calling are detecting contiguous regions of the genome with the same ploidy, called segments, and determining the absolute copy number, or ploidy, of each segment. Previous approaches to calling CNAs using single cells have been based on Hidden Markov Models (HMMs) and change point detection (11). For example, HMMcopy use a Hidden Markov Model to segment tumor genomes using GC and mappability corrected tumor reads, normalized by matched normal cells. Although HMMcopy was originally designed for array comparative genomic hybridization data (12, 13), it’s been widely used for single cell sequencing data (11, 13).

CopyNumber was also designed for microarray use, and uses normalized and log transformed copy number measurements rather than raw read counts to detect breakpoints from changes in genome coverage. However, although this method outputs segments, it does not output absolute copy number calls. One strength of CopyNumber, however, is that it can be run in individual and multi sample modes. In the multi sample mode, breakpoints are forced to be shared across all samples (14).

AneuFinder, which was designed for calling CNAs in single cell whole genome sequencing data, uses a trained HMM to model copy number state using a negative binomial distribution (15). In newer versions, Aneufinder uses change point detection analysis to find changes in read coverage (16). To determine absolute copy number, each segment is normalized and scaled such that the mean bin count matches a known ploidy, which is determined from a DNA quantification technique, such as flow cytometry (17). If overall ploidy is not known, a scalar is fit such that all segments get an integer copy number (15).

Ginkgo uses variably sized bins for GC correction and removes outlier “bad” bins based on a fixed set of diploid cells (18), then employs circular binary segmentation (19) to detect breakpoints in normalized read counts and scales ploidy estimates to call absolute copy number. Ginkgo can also cluster cells and build phylogentic trees (18).

The method SCNV automatically identifies and uses diploid cells as a null error model, and adapts SeqCBS (20) for use in single cells by pooling diploid cells, calibrating model cutoffs using the pooled diploid cells, and discretizing copy number calls (21). SCNV then uses a bin free method based on change point detection on two nonhomogeneous Poisson processes (20) to segment the genome and identify CNAs. This allows for greater resolution of CNAs which might be obscured by choice of bin size boundaries (21).

SCOPE uses a Poisson latent factor model, based on CODEX (22), to normalize read counts, and then uses a log-likelihood ratio test across multiple samples (23) to detect shared breakpoints, with the segmentation stopping rule defined by a cross-sample modified Bayes Information Criterion (24). This allows SCOPE to use cell specific and shared sample information to better estimate technical noise (25).

CHISEL phases SNP haplotypes (26) of fixed size, and uses cell specific read depth ratios and allele specific frequencies to cluster bins across cells in order to call allele specific copy numbers. This allows CHISEL to call CNAs that are aligned with the observed allelic balance, but also makes it prone to errors caused by allelic drop out from low sequencing coverage (27).

SCICoNE corrects read counts for GC and mappability across bins and cells, then uses a likelihood based model to detect breakpoints shared across cells by combining adjacent bins with similar copy number states. SCICoNE then builds a CNA based tree without the infinite sites assumption, allowing for an arbitrary number of CNAs at a site (28). However, the CNA calling procedure precedes and is independent of the tree reconstruction (29).

All of these methods, except for SCNV, require dividing the reference genome into adjacent bins of variable or uniform size. All methods use bin or cell specific GC and mappability corrections to adjust read counts and mask out “bad” bins that exhibit extremely high or low coverage due to centromeres, telomeres, or highly repetitive regions. However, only SCNV and SCOPE utilize detailed bin specific coverage information from diploid cells, and none are based on explicit stochastic models of tumor evolution. An objective of this paper is to develop models for CNA calling based on explicit models of tumor evolution. The rationale is that the use of such explicit models of evolution might improve inferences similarly to what has been observed in models of molecular evolution used in phylogenetics (30–32).

Because tumor cells evolve forward in time from an ancestral diploid state through mutations that only depend on the current state of the cell, copy number alterations are inherently governed by a (possibly time-inhomogenous) temporal Markov process. However, the read distribution observed along the length of the genome (the spatial process) is not Markovian. To realize this, consider a mutation within a segment of DNA with ploidy 4 that reduces the ploidy from 4 to 3. When moving from the left to the right along the length of the genome, the ploidy would then go from 4->3->4. There are two transitions (breakpoints) caused by the same single CNA. In many other situations, the rate of mutation from 3->4 (as in the second breakpoint) might be low, however, because the chromosome previously was in state 4, the rate of transition back from 3 to 4 is in fact high in our example. The process along the length of the genome is not Markovian because copy number alterations may have finite length and each mutation may induce two breakpoints.

Even though the spatial process is not Markovian, the HMM framework is computationally convenient. An aim of this paper is, therefore, to develop Markovian approximations of the spatial process that can be used for inference. We present SCONCE (Single Cell cOpy Numbers in CancEr), a method based on modeling the temporal Markovian evolutionary process and deriving a best approximating spatial HMM from this process. SCONCE also uses diploid data as a null to model the technical noise in single cell sequencing data and can robustly learn model parameters and detect copy number alterations. We show on simulated data that the method more accurately estimates the ploidy states of a cell than previous state-of-the-art methods, and we analyze real data to show that the observations from simulated data are mirrored by similar differences among methods in analyses of real data.

## 2. Theory and Methods

### 2.1. Simulations

In order to robustly evaluate SCONCE, we provide two simulation models, one based on line segments and one based on bins. In particular, the line segment model simulates the evolutionary process behind CNAs without assuming any bins, but treating the genome as a line segment. The binned model divides the genome into discrete bins when simulating the evolutionary process. We consider the line segment process to be the more realistic evolutionary model. Of note, the provided simulation models are derived differently from the Hidden Markov Model, described in Section 2.3. We simulate data and estimate parameters and copy number calls under different models, to avoid biasing method comparisons towards our method. See Supplement S1 for full simulation details.

### 2.2. Simulation Datasets

We simulated 4 datasets under the line segment model and 6 datasets under the binned model in order to generate a variety of types and quantity of copy number events. Specifically, each dataset had 100 tumor cells and 100 diploid cells, where read counts from diploid cells were averaged together to form the null model. See Supplement S1.4 for full simulation parameter values.

### 2.3. Hidden Markov Model

In order to simultaneously segment the tumor genome and call absolute copy numbers, we use a Hidden Markov Model along the length of the genome. We define the state space of the HMM as the integer tumor ploidy in a given genomic bin, from 0 up to a user specified *k* (suggested *k* = 10), and the alphabet as the integer observed tumor read depth in that bin.

We model emission probabilities for tumor read counts per bin with a negative binomial distribution (interpreted here as an overdispersed Poisson). We incorporate the mean diploid read count for each bin into the emission probabilities, in order to normalize for technical noise and sequencing bias. Let the tumor read depth in window *i* for tumor cell *A* be represented by random variable *X_iA_*, such that

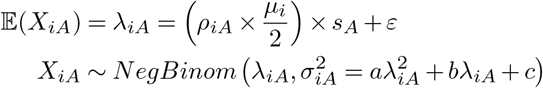

where *ρ_iA_* is the ploidy in window *i* for cell *A*, *μ_i_* is the mean diploid read depth in window *i*, *ε* is a constant sequencing error term, *s_A_* is a cell specific library size scaling factor, and {*a, b, c*} are constants learned from diploid data, such that the emission probability for an observed read depth, *x_iA_*, is given by the specified negative binomial distribution. See Library Size Scaling Factors for *s_A_* calculations and Negative Binomial Mean and Variance Calculations for {*a, b, c*} calculations.

For the HHM, the initial probability vector is defined as the steady state distribution of the Markov chain. The log-likelihood of the observed tumor data is calculated using the forward algorithm and summed across all chromosomes for a given cell. The HMM is reset to the initial probability vector at the beginning of each chromosome to maintain chromosomal independence.

### 2.4. Joint evolutionary process process of two bins forward in time

In Supplement S1, we described two principled models of CNA evolution. However, neither of these models have the property that they are Markovian along the length of the genome. To construct an approximating process that is Markovian, we will first construct a process affecting two bins. This process will effectively be similar to the described binned process, but it is parameterized slightly differently out of convenience. From this description of the joint evolution of two bins, we will then derive the approximating Markov process used for HMM inference of copy number state.

Consider two adjacent bins in the genome on one lineage, (*U, V*) ∈ {(0,0), (0,1),…, (*k,k*)}, where *U* is the ploidy in bin *i*, and *V* is the ploidy in bin *i* + 1. The ploidies in these bins change through evolutionary history according to rate parameters {*α,β,γ*}:

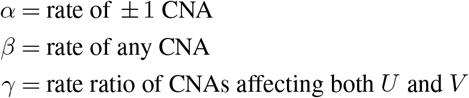

These rates are encoded in a transition rate matrix 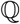:

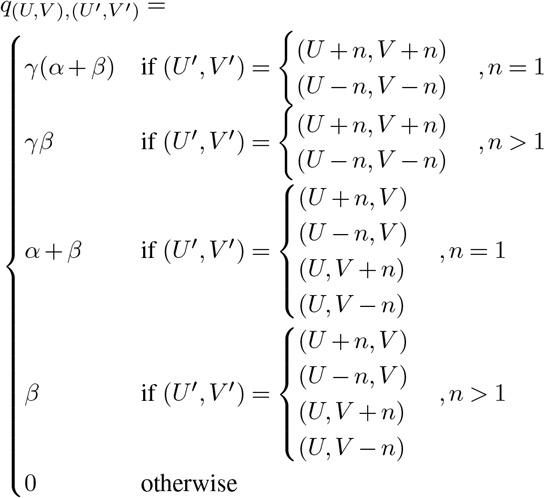

From this rate matrix 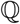, the time dependent transition probabilities 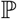 are calculated via the matrix exponential as

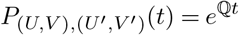

This gives the probability of observing a transition from (*U, V*) to (*U*′, *V*′) in time *t*.

### 2.5. Discrete process (Markovian approximation) along the genome

We convert the forward-in-time process for two bins into a Markov model along the length of the genome with transition probability matrix 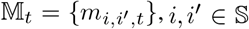, i.e. we identify the probability of moving from state *i* to *i*′ along the genome, after a given evolutionary time *t*. Under the assumption that the cell has an ancestral diploid state at time *t* = 0, we set (*U,V*) = (2,2) and (*U*′, *V*′) = (*i*, *i*′). By normalizing over all states *W* in 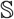, the one-step transition probabilities of the discrete approximating Markov process along the length of the genome are given by

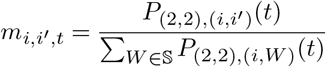

This time dependent transition matrix approximates a non-Markovian process using an evolutionary time-informed HMM. The advantage of using this model over more generic HMMs is that information about the ancestral diploid state is included in the model specification allowing more accurate inference of ploidy state. While the model is only an approximation, as it ignores the non-Markovian nature of any realistic model of CNA changes along the genome, we will evaluate it using the aforementioned non-Markovian simulation models.

### 2.6. Model Training

The model training has four steps. We first estimate the constants, {*a, b, c*}, used to model the emission probabilities, from the diploid data. Second, for each tumor cell, *A*, we quickly estimate an unconstrained transition matrix, initial probability vector, and library size scaling factor, *s_A_*, using a modification of the Baum Welch algorithm. Third, the model rate parameters, {*α_A_,β_A_,γ_A_,t_A_*}, are then fit to the estimated transition matrix using least squares. Fourth, the initial estimates for {*s_A_,α_A_,β_A_,γ_A_,t_A_*} are refined using the Broyden–Fletcher–Goldfarb–Shanno (BFGS) optimization algorithm to maximize the forward loglikelihood of the observed tumor read depths. See Supplement S2 for full model training details.

### 2.7. Real Data preprocessing

We applied SCONCE to a published dataset, consisting of 34 diploid cells (as determined by cell sorting), and 4 tumor subpopulations (24, 24, 4, and 8 cells, respectively) from one triple negative breast cancer patient (33), a cancer type with prevalent CNAs (34).

We first applied standard preprocessing and quality control steps to the sequencing data: trimming adapters and low quality read ends (35, 36), removing low complexity and short reads (37), and removing PCR duplicates (38). After cleaning up the sequencing data, reads were aligned to the reference genome (hg19) using bowtie2 (39), and reads with q scores less than 20 were removed (40).

The reference genome was binned into uniformly sized bins, and cell specific read depth was counted in each dataset using bedtools (41). Per window read depth was averaged across diploid cells, and the {*a, b, c*} constants for the Negative Binomial distribution were calculated (see Supplement S2.1 for full details).

### 2.8. Other methods

In order to evaluate the accuracy of the inference procedure, we compared to HMMcopy (12, 13), CopyNumber (14), and AneuFinder (15, 16). We limited our comparison to methods that do not require bam files or SNPs, as our simulation model does not create bam files or model SNPs for simplicity. For both real and simulated datasets, we used the averaged diploid cells as the matched normal sample to determine the somatic copy number for each tumor cell. To run HMMcopy, the `HMMsegment` function (default parameters) was used to segment each cell, and copy numbers were extracted from the resulting `state` element −1.

The normalized and log-transformed copy number estimates from HMMcopy were used as input for CopyNumber. Then, missing data were imputed, using the `constant` method, and the `winsorize` function was used to remove outliers. To run in single sample mode, the `pcf` function was used with parameters `return.est=T`, `normalize=T`, `digits=6`, and the exponentiated `estimates` element was extracted for the copy number estimates. To run in multi sample mode, the `multipcf` function was run with parameters `return.est=T`, `digits=6`, and copy numbers were similarly extracted from the exponentiated estimates result. Additionally, because CopyNumber does not output absolute copy number calls, we scaled Copy-Number results to minimize the sum of squared differences from the true ploidy in simulated datasets to create a ploidy estimate for comparison purposes.

The procedure for running AneuFinder differed slightly between simulations and real data. See Scripts to run other methods for full scripts.

#### 2.8.1. Simulations

Because our simulation model does not incorporate GC or mappability into simulated read depth, we did not use the GC and mappability corrections in HMM-copy (and subsequently CopyNumber) in order to avoid over-correcting. We used the averaged diploid read counts for the matched normal sample to detect somatic CNAs only. Similarly, in AneuFinder, we skipped the GC and mappability corrections by running the `findCNVs` function with default parameters (`method=“edivisive”, R=10, sig.lvl=0.1`), and extracting the `copy.number` element from the model segments.

#### 2.8.2. Real Data

With real data, we ran HMMcopy by first doing read correction (`correctReadcount`, default parameters) on both the tumor data and averaged diploid cells, then ran as described above.

To run AneuFinder, we ran the `Aneufinder` function with 250,000 binsize, all chromosomes, GC correction, and hg19 assembly. As in simulations, copy number calls were extracted from the `copy.number` element from the `edivisive` model segments.

## 3. Results

### 3.1. GC content and mappability

Because GC content and sequence mappability can bias read distributions, many methods explicitly incorporate corrections for GC content and sequence mappability. However, any technical noise that would affect the tumor sequencing would also affect the diploid sequencing, so in SCONCE, these corrections are already directly accounted for in our emission probabilities via the diploid mean.

To verify this, we examined prediction accuracy of expected tumor read counts per window with different amounts of information. For window *i*, let *μ_i_* be the mean diploid read count, *ζ_i_* be the GC content, and *η_i_* be the mappability from the Duke Uniqueness of 35bp Windows from ENCODE/OpenChrom (UCSC accession wgEncodeEH000325) (42, 43). For each tumor cell, *A*, from the previously published data in (33), we predicted the *i*th window tumor read depth, *x_iA_*, using various linear regressions on {*μ_i_,ζ_i_,η_i_*}, then calculated the sum of squared differences between predicted and actual tumor read depths. Boxplots of the summed squared differences per cell are shown in Figure 1 and empirical cumulative distribution function (ECDF) plots are shown in Supplemental Figure S1 for A: *x_iA_* ~ *μ_i_*, B: *x_iA_* ~ *μ_i_* + *ζ_i_*, C: *x_iA_* ~ *μ_i_* + *η_i_*, D: *X_iA_* ~ *μ_i_* + *ζ_i_* + *η_i_*, E: *x_iA_* ~ *ζ_i_*, F: *x_iA_* ~ *η_i_*, G: *x_iA_* ~ *ζ_i_* + *η_i_*,.

**Figure 1:**
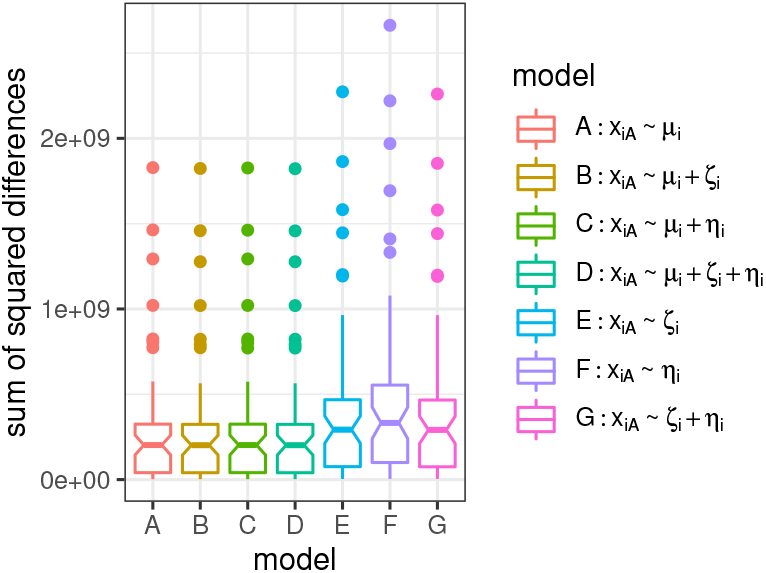
For each linear regression, a boxplot of the sum of squared differences between the predicted read count and observed read count for each tumor cell in (33) (uniformly sized 250kb bins) is shown. No statistically significant difference in error is observed by adding GC or mappability information to the diploid null model.

The sum of squared differences remains consistent across models that incorporate the diploid mean (models A, B, C, and D), and have overlapping ECDF plots, while the sum of squared differences increases for models that depend solely on GC content and mappability (models E, F, and G). Because adding the GC content and mappability did not perform significantly differently from the diploid mean alone (two sample KS-test on the cumulative distribution of summed squared differences, *D* = 0.033333, *p*-value = 1), we conclude using the diploid mean is sufficient, and do not add GC or mappability corrections. This conclusion is robust to changes in window size and binning method (ie. uniformly sized bins vs variably sized bins with equal numbers of uniquely mappable bases).

### 3.2. Error rates

To compare the accuracy of each copy number calling method, we compared the sum of squared differences (SSD) between true copy number and estimated copy number across ten simulation datasets. Recall that these datasets were simulated under a more general non-Markovian model (see Supplement S1 for simulation details and Supplement S1.4 for parameter values).

For each cell, the SSD was calculated across all 12,397 windows (number of uniform non-overlapping 250kb windows in hg19). Overall, SCONCE has similar or lower error rates than AneuFinder, and consistently significantly lower error rate than HMMcopy and CopyNumber (see Figure 2).

**Figure 2:**
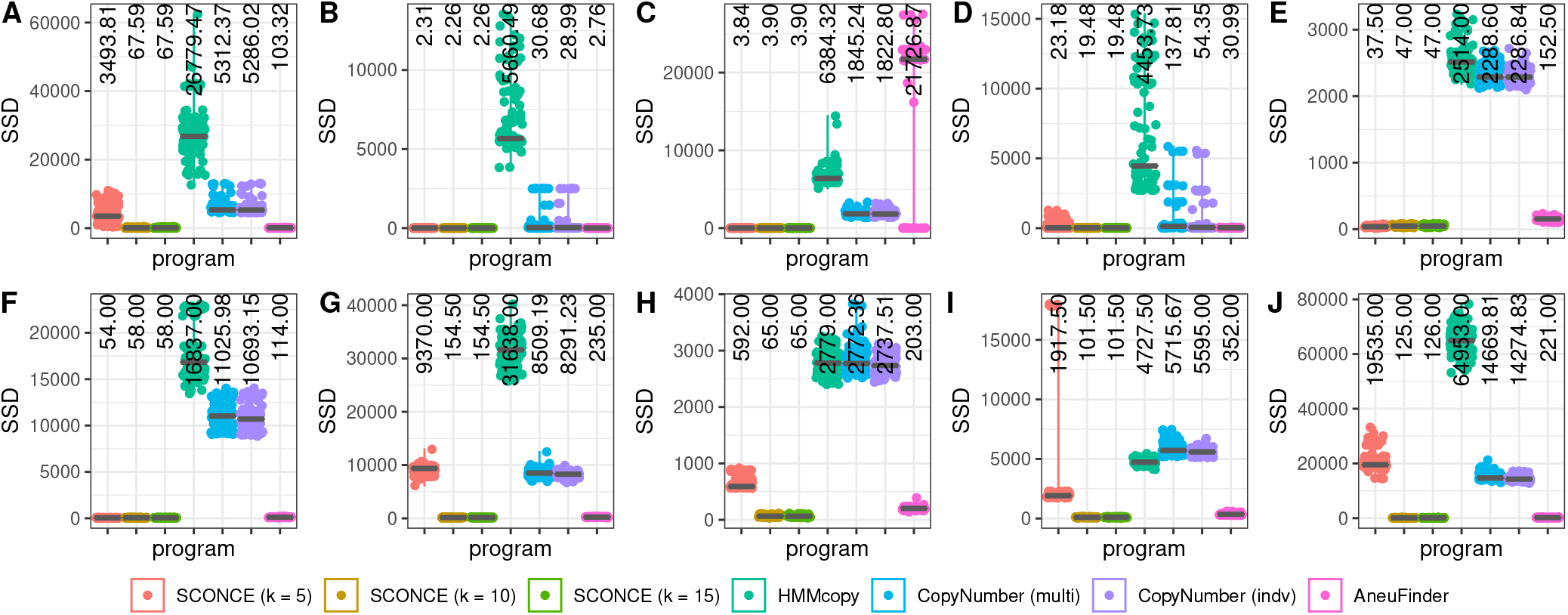
For each method, the sum of squared differences (SSD) between simulated ploidy and estimated ploidy is shown across different parameter sets. Each dot represents the error for one cell and the median SSD is shown with a gray line and printed at the top of each column. SCONCE consistently has SSD values that are lower or on par with other methods.

For example, in Simulation Set A (consisting of many small overlapping CNAs per cell, under the line segment model; Figure 2A), the median SSD for SCONCE is 3493.41 and 67.59, for *k* = 5 and *k* = 10, 15, respectively, which is lower than the median SSD for AneuFinder, at 103.32. Meanwhile, the median SSD for CopyNumber (in multisample and individual modes, respectively) was 5312.37 and 5286.02, and the median SSD for HMMcopy (which does not output absolute copy number calls, and so was optimally scaled) was 26779.47. Of note, because SCONCE cannot call ploidies above the user specified *k*, its error rate is significantly higher for *k* = 5 when the true simulated ploidy is greater than 5.

Scaling problems can also arise if *k* is set too low. For Simulation Set I (consisting of very short spiky CNAs, under the binned model; Figure 2I), the median SSD for SCONCE for *k* = 5 is 1917.50, while the median SSD for *k* = 10, 15 drops to 101.50. The median SSD for AneuFinder is over three times worse at 352.00, while the median SSDs for HMMcopy and CopyNumber (multisample and individual modes) are orders of magnitude worse, at 4727.50, 5715.67, and 5595.00, respectively. In both of these simulation sets, despite the higher median SSD for SCONCE at *k* = 5, the median SSD consistently drops for *k* = 10, 15. Because higher values of *k* result in a higher run times without significant gain in accuracy, we recommend setting *k* = 10.

In other simulations, AneuFinder has scaling problems that SCONCE does not. In Simulation Set C (consisting of mainly deletions, under the line segment model; Figure 2C), the median SSD for SCONCE is 3.84 and 3.9 for *k* = 5 and *k* = 10, 15. The median SSD was 6384.32, 1845.24, and 1822.80 for HMMcopy and CopyNumber (multisample and individual modes), respectively. However, the median SSD for AneuFinder is orders of magnitude higher, at 21726.87. Upon closer inspection, AneuFinder incorrectly doubles the ploidy for the majority of the cells in this simulation set (see Supplemental Figure S3C for an example decoding and Supplemental Figure S2C).

To check if the differences in median SSD between methods were due to scaling issues, we also rescaled all copy number calls to minimize the SSD between simulated ploidy and estimated ploidy for all methods. With this optimal rescaling, SCONCE consistently outperforms or is on par with other methods (see Supplemental Figure S2).

Although the median SSD for SCONCE with *k* = 5 in Simulation Set A decreases from 3493.81 to 2900.76, rescaling does not address the underlying limitation of *k* being too small. The median SSDs for the other methods for Simulation Set A also decrease, but not significantly (see Supplemental Figure S2A). Similarly, under Simulation Set I, fixing the incorrect scaling for SCONCE with *k* = 5 causes the median SSD to drop from 1917.50 to 1649.83, but it doesn’t address the root problem of *k* limiting the ploidies SCONCE can call.

In contrast, the median SSD for AneuFinder for Simulation Set C drops significantly from 21726.87 to 4.33, while the median SSD for SCONCE remained constant. This shows AneuFinder’s high median SSD for Simulation Set C was due to incorrect scaling, rather than incorrect breakpoint detection and segmentation.

However, although HMMcopy median SSD values decreased with optimal scaling, they never dropped into the same range as SCONCE and AneuFinder, implying there are non-scaling related reasons behind the high median SSD values. The median SSD values for CopyNumber did not change, as its output was already scaled because it does not report absolute ploidies.

### 3.3. Genome wide decodings

By plotting the genome wide copy number profile for a representative cell from each simulation set, we can learn more about the specific differences between methods that lead to differing error rates. Of note, the value of *k* must be set high enough to allow a wide enough ploidy range in SCONCE (suggested *k* = 10). For brevity, only genome decodings for SCONCE (with *k* = 10) and AneuFinder are shown in the main text (see Supplemental Figure S3 for decodings with other programs and other values of *k* for SCONCE).

In some cases where the maximum *k* is set too low, the error rate from SCONCE is high because it can’t estimate high enough ploidies (Simulation Set A, with many small overlapping CNAs; Figure 2A, Supplemental Figure S3A). This can be seen in chromosome 3, where the true ploidy reaches a maximum of 8, but SCONCE’s ploidy estimates are limited to *k*.

In other cases, setting *k* too low causes the library size scaling factor to be estimated incorrectly (Simulation Set I, with very short spiky CNAs, Figure 2I, 3A, Supplemental Figure S3B). Specifically, for *k* = 5, SCONCE incorrectly reports ploidy of 1 instead of 2 for most of the genome. However, once the value of *k* is high enough (*k* = 10, 15), SCONCE consistently recovers the simulated ploidy with a lower error rate than AneuFinder. In particular, AneuFinder misses small CNAs (ranging from 1 to 5 bins in width), that SCONCE does not miss, such as in chromosomes 9 and 17 (shown with arrows in Figure 3A). These results are consistent across simulations with approximately equal rates of insertions and deletions (Supplemental Figure S3A, S3B) and in simulations with mostly insertions (Supplemental Figure S3D).

**Figure 3:**
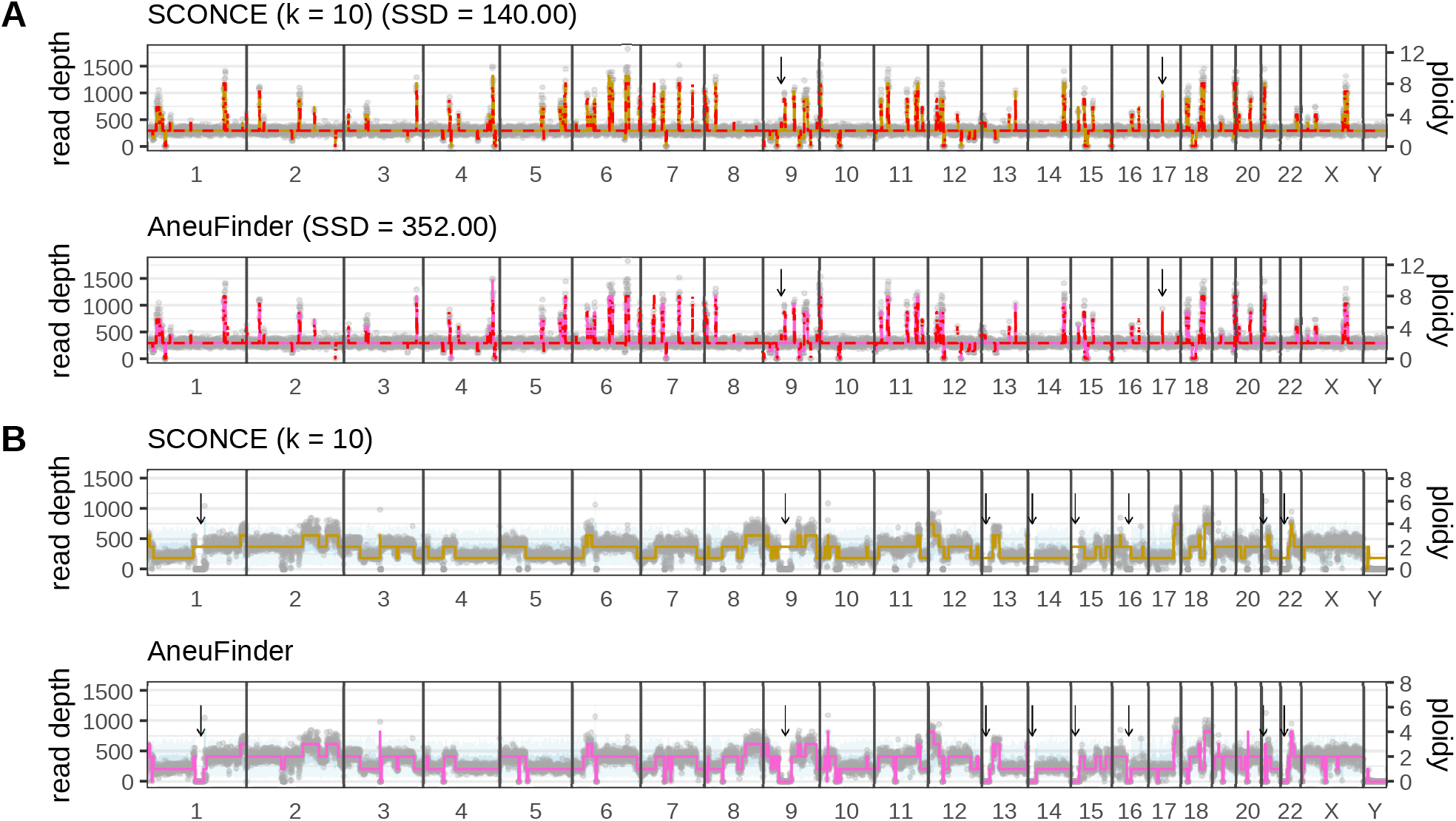
Genome wide copy number decodings are shown for representative cells from simulations and real data. Simulation Set I (very short spiky CNAs under the binned simulation model) is shown in panel A, and cell SRR054570 from (33) is shown in panel B. Genomic window is plotted along the x-axis, per window read depth is shown along the left y-axis, and ploidy is plotted along the right y-axis. Black vertical lines denote chromosome boundaries, gray dots represent observed tumor read depth in each window, the red dotted line denotes the true ploidy from simulation (where applicable), the light blue line shows the mean diploid read count, the light blue band shows ±1 standard deviation in the diploid read count, and the colored lines denote the copy number decoding from each method. Black arrows highlight regions with differences in CNA calls between SCONCE and AneuFinder. In panel A, small CNAs in chromosomes 9 and 17 are called by SCONCE, but not by AneuFinder. In panel B, centromeres in chromosomes 1, 9, and 16 and telomeres in chromosomes 13-15, 21, and 22 are called with ploidy 0 by AneuFinder, but SCONCE makes the more parsimonious call of ploidy 2.

Additionally, in simulations with mostly deletions (Simulation Set C, under the line segment model), AneuFinder consistently and incorrectly doubles the estimated ploidy, leading to a high error rate, while SCONCE does not (Figure 2C, Supplemental Figure S3C). Specifically, AneuFinder mainly calls ploidies of {0, 2, 4}, instead of {0, 1, 2}. When we optimally scaled all copy number estimates to minimize the SSD, AneuFinder’s error rates dropped, thereby verifying the existence of a scaling problem. Even with this optimal scaling, SCONCE continued to have lower error rates than other methods (Supplemental Figure S2).

Furthermore, SCONCE considerably outperforms methods like HMMcopy and CopyNumber in regions of 0 read coverage. By using the diploid null model, we are able to separate between true deletions and areas that have missing data due to sequencing noise (Simulation Set A, with many small overlapping CNAs, Supplemental Figure S3A; Simulation Set C, with mostly deletions, Supplemental Figure S3C). We note that this problem observed in the real data was not contributing to the performance of HMMcopy and CopyNumber in the simulated data, as no regions with missing data were simulated.

For real data (33), copy number estimates from a representative cell (SRR054570) from SCONCE (with *k* = 10) and AneuFinder are shown in Figure 3B (see Supplemental Figure S4 for copy number estimates from each method for cell SRR054570 and another representative cell, SRR053675). Of note, because we specifically incorporate diploid data as our null model, SCONCE makes the most parsimonious calls, rather than assuming copy number 0, in regions that are hard to sequence or map and have no diploid data. For example, in regions around centromeres and telomeres, AneuFinder often calls 0 ploidy when there’s no observed diploid or tumor reads. However, SCONCE uses the lack of diploid and tumor reads to predict no change in copy number. This can be clearly seen in Figure 3B in the centromeres of chromosmes 1, 9, and 16, and in the telomeres of chromosomes 13, 14, 15, 21, and 22. Additionally, by examining Supplemental Figure S4B, small CNAs (between 5 and 22 250kb windows in length, on chromosomes 9, 10, 12, 13, and 18) are consistently missed by AneuFinder, while SCONCE calls these. CNAs larger than 99 windows in length, however, are consistently called well by both SCONCE and AneuFinder. These results recapitulate the results seen from simulations.

## 4. Discussion

CNAs are an important driver in cancer evolution, and accurately detecting them on a single cell level can deepen our understanding of tumorigenesis. In this paper, we derive several models of copy number alterations for inference and simulation. We show that using HMMs derived from models of the evolutionary process that generate CNAs, more accurate inferences of CNA could be obtained. The method for inference based on these models, SCONCE, is available as an open source computer package at https://github.com/NielsenBerkeleyLab/sconce.

One limitation of SCONCE is that it requires data from diploid cells sequenced on the same platform as the tumor cells. While this increases accuracy by accounting for platform specific biases and single cell sequencing errors, it also increases sequencing costs to sequence diploid cells, which may not be directly of interest to investigators. Alternatively, as in the (33) dataset and in other methods (27) utilizing other datasets, sequenced tumor cells that are determined to be diploid by other means (such as via cell sorting) can be relabeled as diploid cells.

One of the key strengths of SCONCE over competing methods is its principled Markovian approximation of a non-Markovian process. This allows for future interpretations and applications of model parameters to understand tumor evolution. Specifically, SCONCE learns transition rate parameters {*α,β,γ*}, tree branch length *t*, and library size scaling factors, but these values are not used directly outside of the copy number profile decoding. Understanding these transition rates in the context of using these tree branch lengths to build phylogenies is the subject of future work.

Compared to other methods, SCONCE has increased sensitivity in calling very small CNAs, particularly those smaller than 5500kb. Additionally, in cells with substantial copy number losses, SCONCE can accurately create copy number profiles without erroneous ploidy doublings. This is due to SCONCE’s method of estimating library sizes using the Viterbi decoding to account for how changes in the copy number profile necessarily impact the library scaling factor.

Furthermore, because SCONCE uses the averaged diploid data as a null model, in regions with zero tumor read coverage, it can differentiate between genomic loss and sequencing noise, which other methods can not do. In particular, in regions with diploid coverage but no tumor reads, SCONCE calls 0 ploidy, and in regions without coverage in either the diploid cells or the tumor cell, SCONCE makes the most parsimonious call. This increases CNA calling accuracy of hard to sequence regions, such as telomeres, centromeres, and repetitive regions.

In conclusion, we present an accurate and principled evolutionary model for calling copy number alterations in single cell whole genome sequencing of tumors, with implications for broader applications.

## 5. Acknowledgements

## 5.1. Funding

This work was supported by the National Institutes of Health [R01GM138634-01 to R.N.].

## 6. Code Availability

SCONCE is implemented in C++11 and is freely available from https://github.com/NielsenBerkeleyLab/sconce. See Supplement S6 for full details.

## Notes

https://github.com/NielsenBerkeleyLab/sconce/

### Competing Interest Statement

The authors have declared no competing interest.

