## Supplement for "SCONCE: A method for profiling Copy Number Alterations in Cancer Evolution using Single Cell Whole Genome Sequencing"

### S1. Simulations

We provide two simulation models, one based on line segments and one based on bins. The line segment model treats the genome as a line segment to simulate the evolutionary process behind CNAs without assuming any bins, while the binned model divides the genome into discrete bins.

#### S1.1. Line segment model.

***S1.1.1. Definitions:.*** We will assume a genome to have a fixed maximal length, $L$, and be comprised of line segments with positions that can be mapped back into $[0,L]$. Each genome may consist of multiple orthologous chromosomes. A chromosome is an ordered list of line segments, $C = \big((b_1,e_1),(b_2,e_2),...,(b_g,e_g)\big)$, $g \in \mathbb{N}$, where $b_i$ and $e_i$ are, respectively, the beginning and end positions of the $i$th chromosomal segments, $b_i < e_i$, $e_i < b_{i+1}$, $b_1 \geq 0$, $e_g \leq L$. Each genome is a set of such chromosomes $G = \{C_1,C_2,...,C_c\}$, $c \in \mathbb{Z}$. The reference genome in a diploid healthy cell is given by $G_h = \big\{\big((0,L)\big),\big((0,L)\big)\big\}$. The length of a chromosome is $\mid C \mid = \sum_{i=1}^{g}(e_i - b_i)$, and the length of a genome is $\mid G \mid = \sum_{i=1}^{c} \mid C_i \mid$.

***S1.1.2. Deletions:.*** A deletion in a chromosome erases part of, or the entire segment, for one or more line segments in a single chromosome. For example, a deletion in a chromosome of a healthy diploid cell between positions $d \geq 0$ and $f \leq L$ would results in a new genome $G' = \big\{\big((0,d),(f,L)\big),\big((0,L)\big)\big\}$. If the same chromosome next is hit by a deletion between position $i$ and $j$ with $0 \geq i \leq d$ and $f \geq j \leq L$, the new genome would be $G'' = \big\{\big((0,i),(j,L)\big),\big((0,L)\big)\big\}$. If this chromosome is hit by a deletion with start and end positions 0 and $l < d$, the new genome will then be $G''' = \big\{\big((l,d),(f,L)\big),\big((0,L)\big)\big\}$ and so forth.

***S1.1.3. Amplifications:.*** An amplification creates an extra copy of a chromosome or part of a chromosome. For example, an amplification starting and ending at positions $m \geq 0$ and $o \leq L$ for in a healthy cell, $G_h$ would result in a genome of composition $G^{*} = \big\{\big((0,L)\big),\big((0,L)\big)\,\big((m,o)\big)\big\}$. An amplification in the same positions in the first chromosome of genome $G'$ from the previous section would result in $G^{**} = \big\{\big((0,d),(f,L)\big),\big((0,L)\big),\big((m,d),(f,o)\big)\big\}$, if $m < d$ and $o > f$. Note there is no maximum ploidy imposed by this model.

***S1.1.4. Rates of amplification and deletion:.*** It is assumed that amplifications and deletions initiate at a constant rates $\varphi$ and $\delta$, respectively, per unit chromosome and per time unit. We will further assume that the lengths are drawn from a truncated exponential distributions with rates $\tau_a$ and $\tau_d$, for amplifications and deletions respectively. The truncation occurs when an amplification or deletion extends beyond the end of the chromosome and we assume that amplifications and deletions run from left to right. To understand this process we can imagine that initiation points of amplifications or deletions are laid down according to a Poisson process with constant rate along the length of the chromosome. To construct the process such that there are no edge effects and a constant rate of deletion and amplification at any point of the chromosome, we furthermore assume that new amplifications and deletions additionally initiate at the left start of each chromosome at rate $\frac{\varphi}{\tau_a}$ and $\frac{\delta}{\tau_d}$, respectively. This ensure that the rate at which a particular point in the region is affected by an amplification or deletion is $\frac{\varphi}{\tau_a}$ and $\frac{\delta}{\tau_d}$, respectively, no matter what the position is, i.e. there are no edge effects with this construction.

The total genomic rate at which amplifications and deletions occur at any point in time is then $c\left(\frac{\varphi}{\tau_a} + \frac{\delta}{\tau_d}\right) + \mid G \mid (\varphi + \delta)$

***S1.1.5. Induced marginal process:.*** The process, as defined here, is a Markov process with state space on the infinite set of all possible genomes. It also induces a marginal continuous time Markov process at each position in the genome, $W_t \in \mathbb{Z}$, with transition rates

$$q_{ij} = \begin{cases} i\frac{\varphi}{\tau_a} \text{ if } j = i+1 \\ i\frac{\delta}{\tau_d} \text{ if } j = i-1 \text{ and } j \geq 0 \\ 0 \text{ else} \end{cases}$$

We notice that this is a linear birth-death process with birth rate $\frac{\varphi}{\tau_a}$ and death rate $\frac{\delta}{\tau_d}$.

**S1.2. Binned process.** We also consider an alternative and simpler process which we will call the *binned process*. In this process we assume that the genome can be divided into $n$ bins. The state space in each bin is $\mathbb{S} = \{0,1,2,...,k\}$, where $k$ is the maximum ploidy.

***S1.2.1. Amplifications and deletions:.*** We assume that the length of amplifications and deletions follows a truncated geometric distribution with parameter $p$. That is, given that a certain amplification/deletion occurs in bin $i$, the probability that it extends to the right to bin $i+1$ is $1-p$ . However, amplifications and deletions may also initiate to the left of the first bin to ensure an approximately constant rate along the length of the chromosome. If the effect of an amplifications/deletion is to change the



DRAFT

| Simulation set | Short description | Deletion rate, $\frac{\delta}{\tau_d}$ | Insertion rate, $\frac{\varphi}{\tau_a}$ | Mean deletion length, $\tau_d$ | Mean insertion length, $\tau_a$ |
|---|---|---|---|---|---|
| A | many small overlapping CNAs | 0.001 | 0.001 | 10 | 10 |
| B | few large CNAs | 0.0001 | 0.0001 | 10 | 10 |
| C | many deletions | 0.0005 | 0.0001 | 10 | 10 |
| D | many insertions | 0.0001 | 0.0005 | 10 | 10 |

Table S1: Description of simulation sets under the line segment model, relative to a genome length of 100. All simulations were done under the neutral coalescent, with a tree branch length leading to the root of tree (the ancestral diploid genome) of 1.0. Although the mean CNA length remains constant across simulation sets, the overall size of CNAs is also affected by the deletion and insertion rates, as CNAs tend to overlap.

| Simulation set | Short description | Max ploidy, $k$ | Tree branch length, $t$ | Geometric $p$ |
|---|---|---|---|---|
| E | very short spiky CNAs | 5 | 1.0 | 0.1 |
| F | long segments | 5 | 100 | 0.005 |
| G | long segments | 8 | 100 | 0.005 |
| H | very short spiky CNAs | 8 | 100 | 0.1 |
| I | very short spiky CNAs | 8 | 100 | 0.1 |
| J | longer segments | 8 | 100 | 0.005 |

Table S2: Description of simulation sets under the binned model, relative to a genome length of 100. All simulations were done under the neutral coalescent, with a tree branch length leading to the root of tree of 1.0.

state in the first bin (in which the amplification/deletion initiated) by an amount $i$, the effect on another bin affected by the same amplification/deletion, currently in state $j$, is to change the state of this bin to $i+j$ if $0 \leq i+j \leq k$. Otherwise, if $i+j$ is less than 0 or larger than $k$, it transitions to 0 or $k$, respectively.

***S1.2.2. Marginal process of initiation:.*** We model the marginal process of initiation of new deletions/amplifications in each bin as a continuous time Markov chain with rate matrix $Q = \{q_{ij}\}$. The total rate of amplification/deletion initiation within any of the $n$ bins, at time $t$, is then $R_t = \sum_{i=1}^{n} \sum_{j \neq Y_i(t)} q_{Y_i(t)} j$, where $Y_i(t)$ is the state in bin $i$ at time $t$. Notice, that because of the assumption of geometrically distributed lengths of amplifications and deletions, the marginal process in each bin does not follow $Q$. Only the initiation process of new amplifications and deletions follows $Q$.

In addition to the amplifications/deletions initiated in one of the $n$ bins in the chromosome, we also assume that an amplification/deletion can initiate immediately to the left of the first bin. Such events occur at a rate of $\frac{R_t}{np}$ and the transition type is then drawn based on the state in the first bin, i.e. the relative probability of change to state $j$ from state $Y_0(t)$ is given by $q_{Y_0(t)} j$.

DRAFT

**S1.3. Read Depth Simulation.** Both the line segment and binned models simulate observed read depth for a given number of genomic windows directly from the simulated genome, $G$. Specifically, read depths are simulated from a Negative Binomial distribution, where the mean observed read depth for each window is defined such that the expected total number of reads equals a constant, $\xi$, specified by the user. The parameter, $r$, of the negative binomial that controls the variance is also specified by the user. Additionally, the user can specify if read coverage should be uniform in expectation across the genome before accounting for CNAs, or the user can specify the expected distribution of reads across the genome.

**S1.4. Simulation dataset parameter values.** For simulations under both the line segment and binned models, the genome was binned into 12,397 bins to match the number of uniform 250kb bins in hg19, the negative binomial parameter $r$ was set to 50, the total number of expected reads, $\xi$, was set to 4,000,000, and cells were simulated with even coverage in expectation before accounting for CNAs. Simulation parameter values and dataset descriptions for the line segment and binned models are shown in Table S1 and Table S2, respectively. The rate matrices, $Q$, for initializing the amplification and deletion processes for the binned model are shown in Supplement Table S3.

### S2. Model training details

**S2.1. Negative Binomial Mean and Variance Calculations.** Earlier, we defined the random variable, $X_{iA}$, as the observed read depth for tumor cell $A$ in window $i$, with the per window variance, $\sigma^2_{iA}$, of the Negative Binomial distribution modeled as a second degree polynomial of the mean, $\lambda_{iA}$:

$$X_{iA} \sim NegBinom\left(\lambda_{iA}, \sigma^2_{iA} = a\lambda^2_{iA} + b\lambda_{iA} + c\right)$$

To determine the constants $\{a, b, c\}$, let $d_{iA}$ be the read depth in window $i$ for diploid cell $A$, such that

$$\begin{aligned}
D &= \sum_i \sum_A d_{iA} \\
w_i &= \frac{\sum_A d_{iA}}{D} \\
c_A &= \frac{\sum_i d_{iA}}{D} \\
\mathbb{E}(d_{iA}) &= w_i \times c_A \times D \\
\mathbb{E}(\sigma^2_{iA}) &= a\mathbb{E}(d_{iA})^2 + b\mathbb{E}(d_{iA}) + c
\end{aligned}$$

$$\begin{bmatrix}
0.0 & 0.0 & 0.0 & 0.0 & 0.0 & 0.0 \\
0.2 & -1 & 0.2 & 0.2 & 0.2 & 0.2 \\
0.2 & 0.2 & -1 & 0.2 & 0.2 & 0.2 \\
0.2 & 0.2 & 0.2 & -1 & 0.2 & 0.2 \\
0.2 & 0.2 & 0.2 & 0.2 & -1 & 0.2 \\
0.2 & 0.2 & 0.2 & 0.2 & 0.2 & -1
\end{bmatrix}$$

$$\begin{bmatrix}
0.0 & 0.0 & 0.0 & 0.0 & 0.0 & 0.0 \\
0.01 & -1 & 0.39 & 0.1 & 0.3 & 0.2 \\
0.1 & 0.2 & -1 & 0.4 & 0.2 & 0.1 \\
0.1 & 0.2 & 0.25 & -1 & 0.2 & 0.25 \\
0.2 & 0.1 & 0.2 & 0.3 & -1 & 0.2 \\
0.1 & 0.3 & 0.2 & 0.2 & 0.2 & -1
\end{bmatrix}$$

Simulation set E

Simulation set F

$$\begin{bmatrix}
0.0 & 0.0 & 0.0 & 0.0 & 0.0 & 0.0 & 0.0 & 0.0 & 0.0 \\
0.21 & -1 & 0.39 & 0.1 & 0.1 & 0.05 & 0.05 & 0.05 & 0.05 \\
0.1 & 0.3 & -1 & 0.3 & 0.1 & 0.1 & 0.05 & 0.025 & 0.025 \\
0.05 & 0.1 & 0.3 & -1 & 0.25 & 0.2 & 0.05 & 0.025 & 0.025 \\
0.1 & 0.1 & 0.1 & 0.2 & -1 & 0.2 & 0.1 & 0.1 & 0.1 \\
0.05 & 0.1 & 0.15 & 0.2 & 0.3 & -1 & 0.1 & 0.05 & 0.05 \\
0.05 & 0.1 & 0.1 & 0.1 & 0.1 & 0.25 & -1 & 0.2 & 0.1 \\
0.03 & 0.17 & 0.1 & 0.1 & 0.2 & 0.1 & 0.2 & -1 & 0.1 \\
0.0 & 0.07 & 0.13 & 0.1 & 0.2 & 0.1 & 0.2 & 0.2 & -1
\end{bmatrix}$$

$$\begin{bmatrix}
0.0 & 0.0 & 0.0 & 0.0 & 0.0 & 0.0 & 0.0 & 0.0 & 0.0 \\
0.21 & -1 & 0.39 & 0.1 & 0.1 & 0.05 & 0.05 & 0.05 & 0.05 \\
0.1 & 0.3 & -1 & 0.3 & 0.1 & 0.1 & 0.05 & 0.025 & 0.025 \\
0.05 & 0.1 & 0.3 & -1 & 0.25 & 0.2 & 0.05 & 0.025 & 0.025 \\
0.1 & 0.1 & 0.1 & 0.2 & -1 & 0.2 & 0.1 & 0.1 & 0.1 \\
0.05 & 0.1 & 0.15 & 0.2 & 0.3 & -1 & 0.1 & 0.05 & 0.05 \\
0.05 & 0.1 & 0.1 & 0.1 & 0.1 & 0.25 & -1 & 0.2 & 0.1 \\
0.03 & 0.17 & 0.1 & 0.1 & 0.2 & 0.1 & 0.2 & -1 & 0.1 \\
0.0 & 0.07 & 0.13 & 0.1 & 0.2 & 0.1 & 0.2 & 0.2 & -1
\end{bmatrix}$$

Simulation set G

Simulation set H

$$\begin{bmatrix}
0.0 & 0.0 & 0.0 & 0.0 & 0.0 & 0.0 & 0.0 & 0.0 & 0.0 \\
0.125 & -1 & 0.125 & 0.125 & 0.125 & 0.125 & 0.125 & 0.125 & 0.125 \\
0.125 & 0.125 & -1 & 0.125 & 0.125 & 0.125 & 0.125 & 0.125 & 0.125 \\
0.125 & 0.125 & 0.125 & -1 & 0.125 & 0.125 & 0.125 & 0.125 & 0.125 \\
0.125 & 0.125 & 0.125 & 0.125 & -1 & 0.125 & 0.125 & 0.125 & 0.125 \\
0.125 & 0.125 & 0.125 & 0.125 & 0.125 & -1 & 0.125 & 0.125 & 0.125 \\
0.125 & 0.125 & 0.125 & 0.125 & 0.125 & 0.125 & -1 & 0.125 & 0.125 \\
0.125 & 0.125 & 0.125 & 0.125 & 0.125 & 0.125 & 0.125 & -1 & 0.125 \\
0.125 & 0.125 & 0.125 & 0.125 & 0.125 & 0.125 & 0.125 & 0.125 & -1
\end{bmatrix}$$

Simulation set I

$$\begin{bmatrix}
0.0 & 0.0 & 0.0 & 0.0 & 0.0 & 0.0 & 0.0 & 0.0 & 0.0 \\
0.125 & -1 & 0.125 & 0.125 & 0.125 & 0.125 & 0.125 & 0.125 & 0.125 \\
0.125 & 0.125 & -1 & 0.125 & 0.125 & 0.125 & 0.125 & 0.125 & 0.125 \\
0.125 & 0.125 & 0.125 & -1 & 0.125 & 0.125 & 0.125 & 0.125 & 0.125 \\
0.125 & 0.125 & 0.125 & 0.125 & -1 & 0.125 & 0.125 & 0.125 & 0.125 \\
0.125 & 0.125 & 0.125 & 0.125 & 0.125 & -1 & 0.125 & 0.125 & 0.125 \\
0.125 & 0.125 & 0.125 & 0.125 & 0.125 & 0.125 & -1 & 0.125 & 0.125 \\
0.125 & 0.125 & 0.125 & 0.125 & 0.125 & 0.125 & 0.125 & -1 & 0.125 \\
0.125 & 0.125 & 0.125 & 0.125 & 0.125 & 0.125 & 0.125 & 0.125 & -1
\end{bmatrix}$$

Simulation set J

Table S3: Amplification and deletion initialization rate matrices for the binned model simulations are shown here for each parameter set.

DRAFT

where $D$ gives the total number of diploid reads, $w_i$ gives the proportion of total reads in window $i$, $c_A$ gives the proportion of total reads in diploid cell $A$, and $\mathbb{E}(d_{iA})$ and $\mathbb{E}(\sigma^2_{iA})$ give the expected mean and variance of the diploid read counts under this second degree polynomial model, respectively.

By parameterizing a Negative Binomial distribution with mean, $\mathbb{E}(d_{iA})$, and variance, $\mathbb{E}(\sigma^2_{iA})$, we can calculate the log-likelihood of the observed diploid data, for a given set of constants $\{a, b, c\}$. We maximize the loglikelihood using Nelder-Mead method for optimization to find the optimal values of $\{a, b, c\}$ for the diploid data, and these values are then carried over to the tumor emission probability calculations. See Diploid Cell Processing Calculations and Code for the code.

**S2.2. Library Size Scaling Factors.** Because each sequenced cell will have a different total number of reads, the expected number of reads for each cell needs to be scaled accordingly. Notably, we calculate these cell specific library size scaling factors in a way that accounts for changes in the distribution of reads across the genome caused by CNAs. Let $T_A$ = total # reads in tumor cell $A$ (across $n$ windows), and recall for window $i$

$$\begin{aligned}
\mathbb{E}(X_{iA}) &= \left(\rho_{iA} \times \frac{\mu_i}{2}\right) \times s_A + \varepsilon \\
\mathbb{E}(T_A) &= \sum_{i=1}^{n} \left[\left(\rho_{iA} \times \frac{\mu_i}{2}\right) \times s_A + \varepsilon\right] \\
&= n\varepsilon + s_A \sum_{i=1}^{n} \left(\rho_{iA} \times \frac{\mu_i}{2}\right) \\
\hat{s}_A &= \frac{T_A - n\varepsilon}{\sum_{i=1}^{n} \left(\rho_{iA} \times \frac{\mu_i}{2}\right)}
\end{aligned}$$

We define $\rho_{iA}$ as ploidy in the $i$th window from cell $A$'s Viterbi decoding path, updated after each iteration of the Baum Welch algorithm, such that the library size scaling factor estimate continually incorporates changes in estimated ploidy across the genome (see Baum Welch Starting Point).

**S2.3. Baum Welch Starting Point.** We use a modification of the Baum Welch EM algorithm to estimate the transition matrix and initial probability vector. Instead of estimating an emission matrix directly, we estimate the library size scaling factor, which determines the negative binomial distribution for emission probabilities (see Library Size Scaling Factors).

Because the initial estimate of the library size scaling factor, $s_{A,initial}$, would be erroneously estimated from the Viterbi decoding from an untrained HMM, we instead run Baum Welch from three starting estimates of $s_{A,initial}$. Additionally, binning the genome into discrete, non-overlapping bins leads to some copy number alterations being split across bin boundaries, as the CNA breakpoints may not align with the bin boundaries. This leads to bins with fractional ploidy when CNAs are averaged over the entire bin. Because the HMM is restricted to integer copy numbers, in some cases, the HMM incorrectly attempts to fit these fractional bins to integer copy numbers by doubling all ploidies and halving the library size scaling factor, since this has a slightly higher forward loglikelihood than rounding the fractional bins under the true library size scaling factor. This leads to transient bins, where the Viterbi decoding strictly increases or decreases through one state for exactly one bin (for example, observing ploidies 0->1->2 across only 3 bins).

To detect and avoid this library size scaling factor misspecification, we use the presence of these transient bins to disqualify library size scaling estimates. Specifically, we first run Baum Welch three times, with different initial values of $\hat{s}_{A,initial}$. For the first run, we set $s_{A,initial,1} = \frac{\text{total \# reads in cell } A}{\text{average \# reads in diploid cells}}$. For the second and third runs, we set $\hat{s}_{A,initial}$ to $\hat{s}_{A,final,1}$ multiplied by 2 and 4, respectively.

Next, we pick the Baum Welch run and $\hat{s}_{A,final}$ estimate that has the highest forward loglikelihood, unless it has at least five instances of transient states across the genome. If so, it will be discarded in favor of the run with the next best forward loglikelihood, if the second best run has less than five instances of transient states or is less than 100 loglikelihood units worse than the best run. This selection procedure corrects for any incorrect estimates of $s_A$ that would lead to ploides of $\{0, 2, 4\}$ instead of $\{0, 1, 2\}$, for example.

Finally, we fit our model parameters, $\{\alpha_A, \beta_A, \gamma_A, t_A\}$ to the estimated transition matrix using least squares to minimize the sum of squared differences between the Baum Welch estimated transition matrix and the transition matrix determined by the rate parameters.

**S2.4. BFGS Parameter Estimation.** Given estimates from previous steps, the parameters $\{s_A, \alpha_A, \beta_A, \gamma_A, t_A\}$ are refined for each tumor cell $A$, by maximizing the forward loglikelihood of the observed tumor data using the BFGS optimization algorithm. The central difference approximation is used to calculate the gradient.

To account for instability in small numbers due to machine encoding, the loglikelihood in each position in the sequence is scaled by the maximum of the loglikelihoods in that position, and the final loglikelihood is re-scaled appropriately.

DRAFT

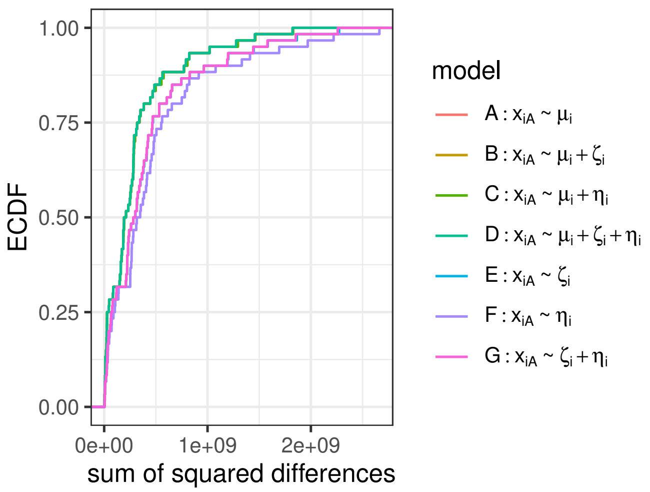


Figure S1: The empirical cumulative distribution function (ECDF) of the sum of squared differences shows no significant differences in adding GC or mappability information. The ECDF lines from models A (diploid mean only), B (diploid mean with GC content), C (diploid mean with mappability), and D (diploid mean with GC and mappability) all lie on top of each other, and the ECDF lines from models E (GC only) and G (GC with mappability) lie on top of each other as well.

Additionally, BFGS performs an unconstrained optimization, but to have sensible results, we require $\{s_A, \alpha_A, \beta_A, \gamma_A, t_A > 0\}$. As such, we log transform and scale $\{s_A, \alpha_A, \beta_A, \gamma_A, t_A\}$ to constrain the optimization results.

Finally, the most likely copy number sequence for each cell is reported using the Viterbi decoding algorithm.

We note that some of the heuristics described in the previous sections could be avoided using a full likelihood estimation using BFGS from the start and avoiding Baum Welch optimization of an approximating unconstrained transition probability matrix. However, we find that such optimization is slower, as the Baum Welch optimization is substantially faster than the BFGS optimization. Furthermore, using only BFGS without any heuristics does not address the inference errors that may occur because the genome is divided into bins that may not be respected by the true CNA mutation process. Finally, using multiple starting points, as described above, was found to be necessary to avoid the optimization to get stuck in local, but not global, optima.

### S3. GC content and mappability

In order to evaluate the information gain by adding GC content and genome mappability, we compared the empirical cumulative distribution functions across several different linear regression models. As shown in Supplemental Figure S1, no significant difference is seen between models based on the diploid mean alone and models utilizing the diploid mean and mappability.

DRAFT

### S4. Scaled Error Plots

As in Figure 2, the sum of squared differences between simulated ploidy and estimated ploidy is shown across different parameter sets for each method. Here, to eliminate scaling errors, all copy number calls are first scaled to minimize the sum of squared differences between simulated ploidy and estimated ploidy. Although the overall SSD values decrease with optimal scaling, SCONCE continues to have SSD values that are lower or on par with other methods.

### S5. Additional Genome Traces

**S5.1. Simulations.** Additional genome wide copy number decodings are shown in Supplemental Figure S3 for representative cells from different simulation conditions, across all methods. Simulation Set A (many small overlapping CNAs under the line segment model) is shown in panel A, Simulation Set I (very short spiky CNAs under the binned simulation model) is shown in panel B, Simulation Set C (mainly deletions under the line segment model) is shown in panel C, Simulation Set D (mainly insertions under the line segment model) is shown in panel D.

**S5.2. Real data.** Additional genome wide copy number decodings are shown in Supplemental Figure S4 across all methods for representative cells SRR054570 (shown in panel A) and SRR053675 (shown in panel B).

### S6. Code Availability

**S6.1. Data preprocessing.** Reads were trimmed using cutadapt (35) and trimmomatic (36), and low complexity reads were removed using prinseq (37). Cleaned reads were aligned to hg19 using bowtie2 (39), reads with a q-score less than 20 were

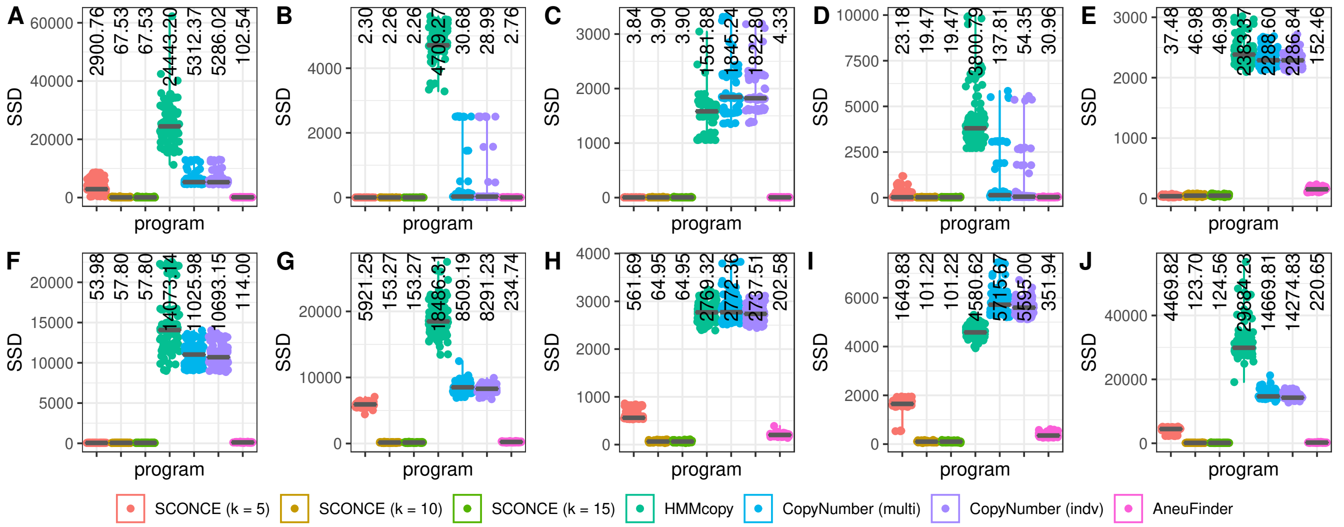


Figure S2: The sum of square differences (SSD) between simulated and estimated ploidies is shown across parameter sets for each CNA calling program. Of note, all estimated ploidies have been scaled to minimize the SSD between simulated and estimated ploidies, in order to eliminate scaling issues in the estimated CNAs.

removed using samtools (40), and duplicates were removed using picard (38). Finally, bedtools was used to create uniform windows of hg19, and to calculate observed read depth per window (41).

**S6.2. Diploid Cell Processing Calculations and Code.** To create the average diploid read count file, the per window read counts were averaged across all diploid cells. An R script (`avgDiploid.R`) is provided to do this.

The variance for the negative binomial distribution on tumor read counts is determined by a second degree polynomial of the mean (see Negative Binomial Mean and Variance Calculations. The calculations to find optimal values of $\{a,b,c\}$ should be rerun for each dataset, using the provided `fitMeanVarRlnshp.R` script. $\{a,b,c\}$ values from one dataset (33) are supplied as defaults.

**S6.3. SCONCE Prerequisites and Dependencies.** SCONCE is implemented in C++11, and requires the Boost C++ Libraries and the GNU Scientific Library (GSL). SCONCE has been extensively tested on Ubuntu 18.04, and is available on github: `https://github.com/NielsenBerkeleyLab/sconce`.

The simulation program for both the line segment and binned models is also available on github. Additional R scripts are provided for intermediate calculations (see Diploid Cell Processing Calculations and Code) and to plot results.

**S6.4. Scripts to run other methods.** Scripts to run HMMCopy, CopyNumber, and AneuFinder as described in Other methods are also provided on GitHub.

DRAFT

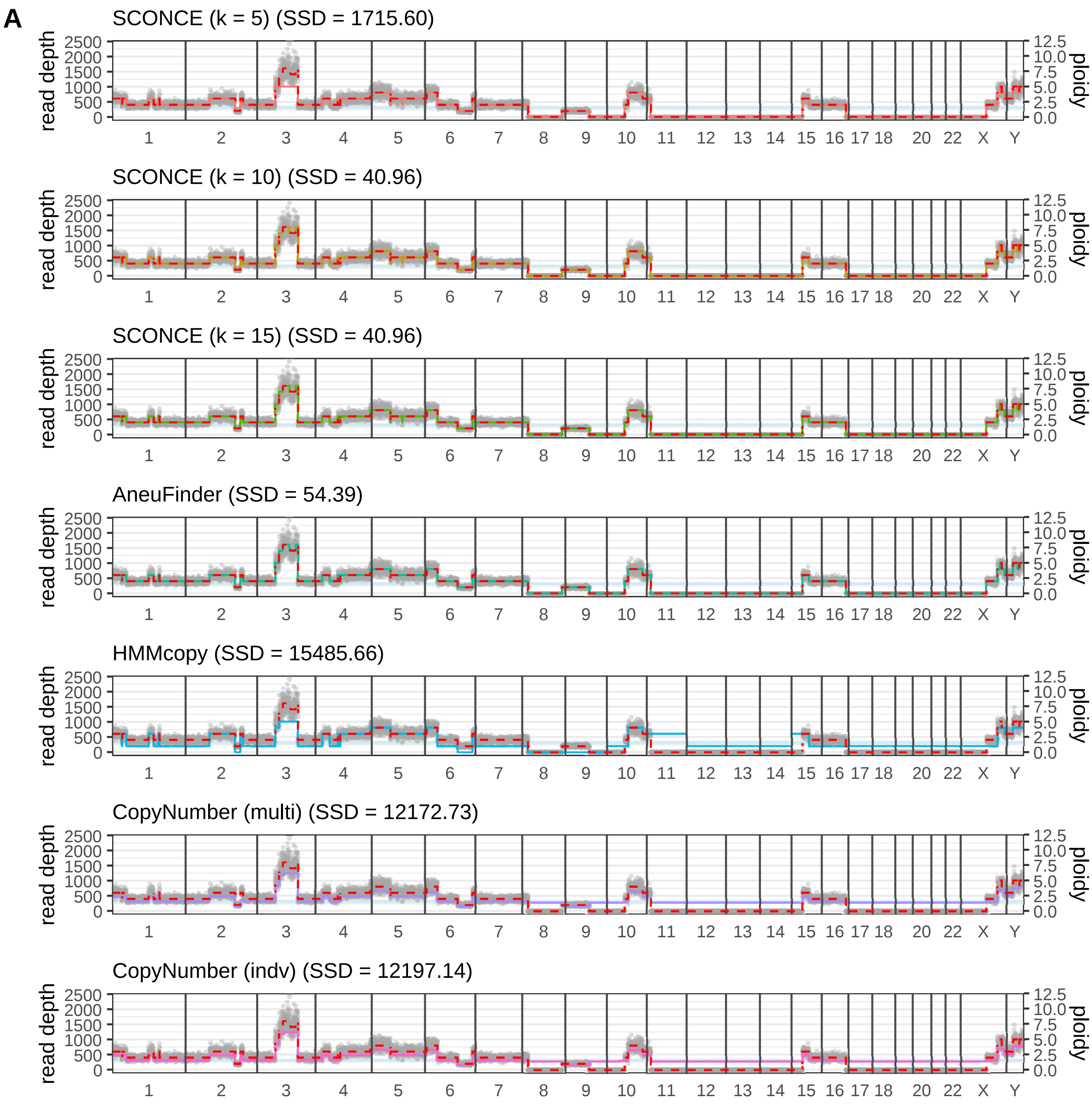


Fig. S3A: Genome wide decoding from cell 0 in Simulation Set A (many small overlapping CNAs under the line segment model).

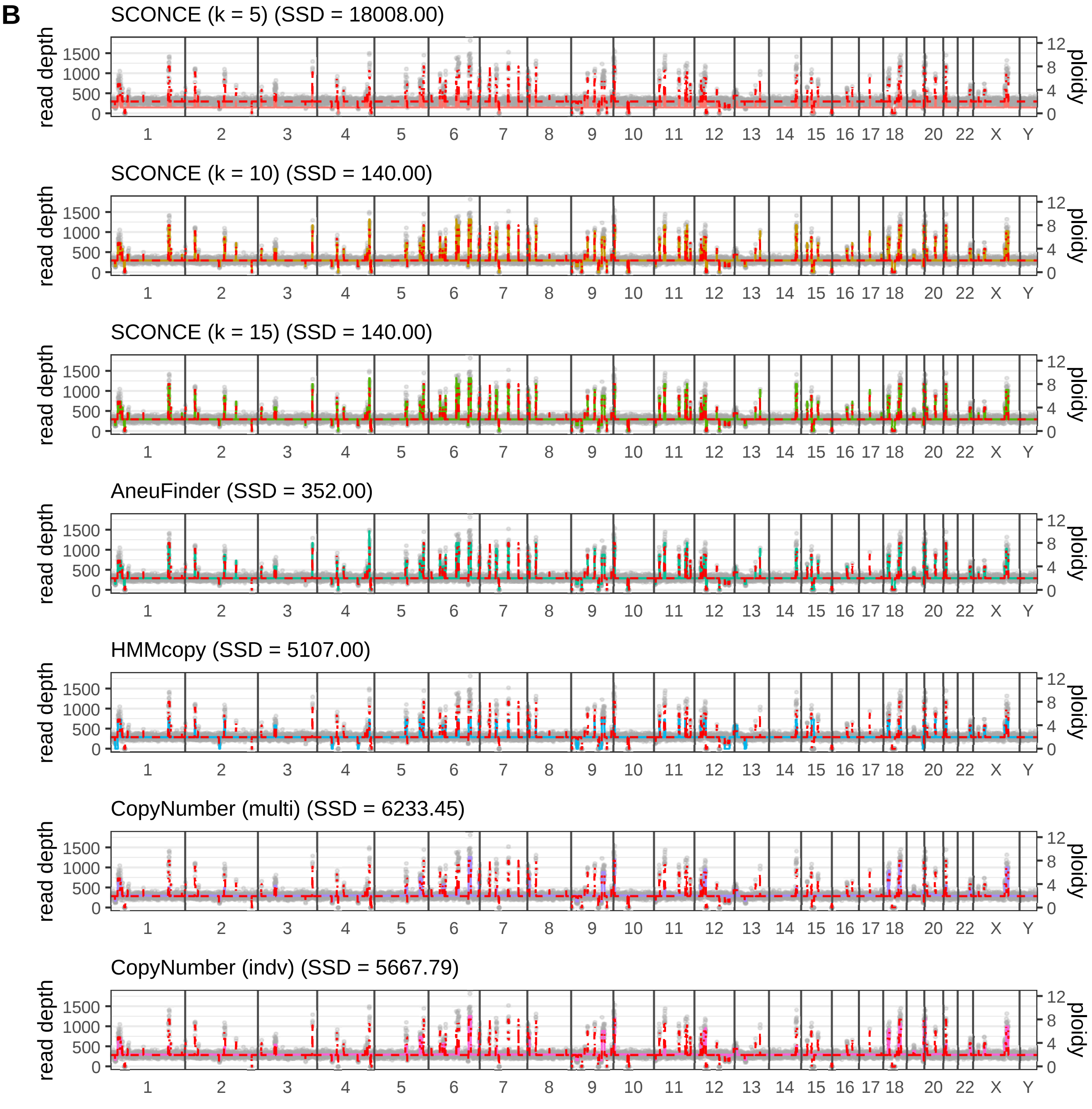


Fig. S3B: Genome wide decoding from cell 54 in Simulation Set I (very short spiky CNAs under the binned simulation model).

**C**

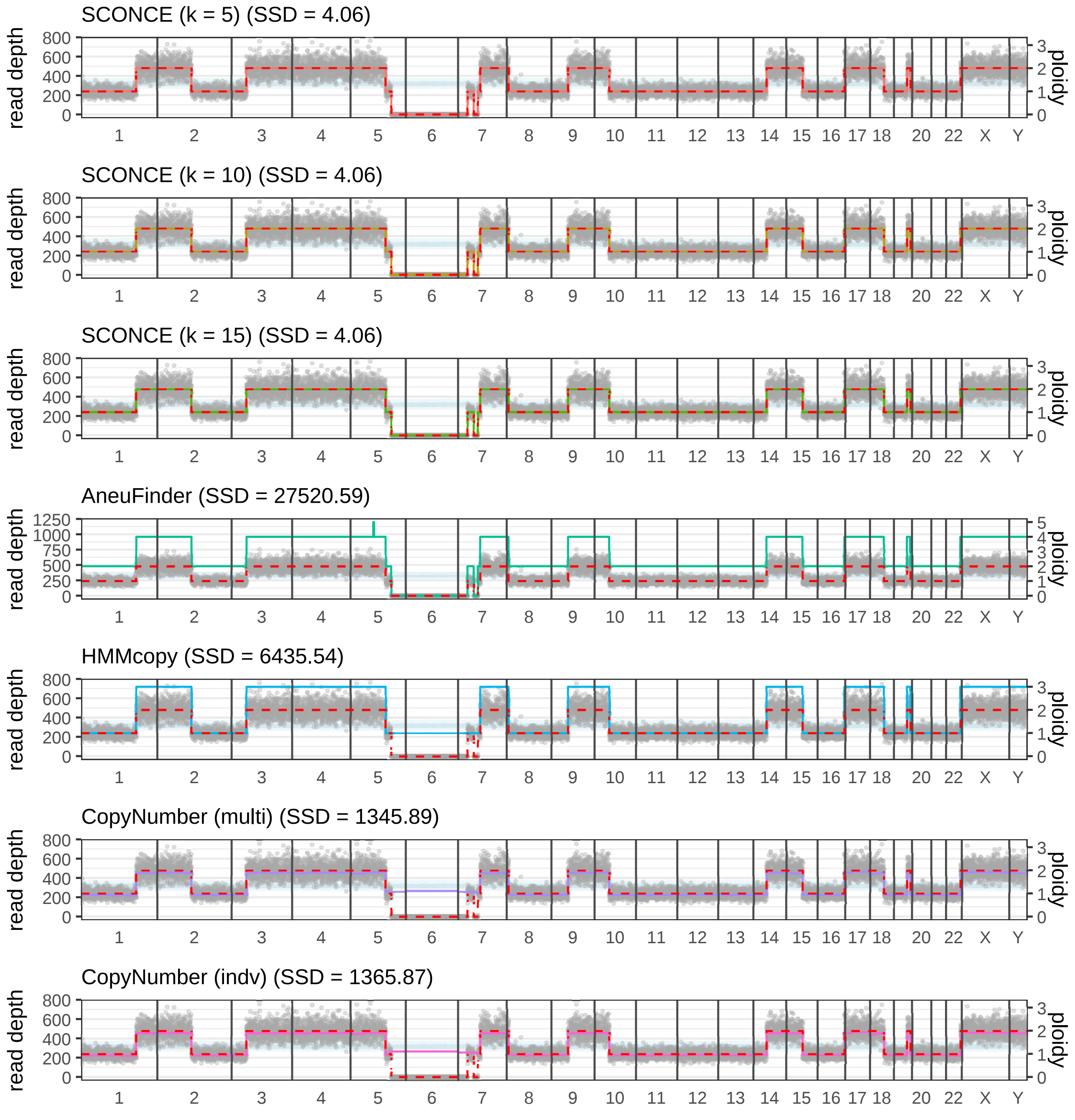


Fig. S3C: Genome wide decoding from cell 14 in Simulation Set C (mainly deletions under the line segment model).

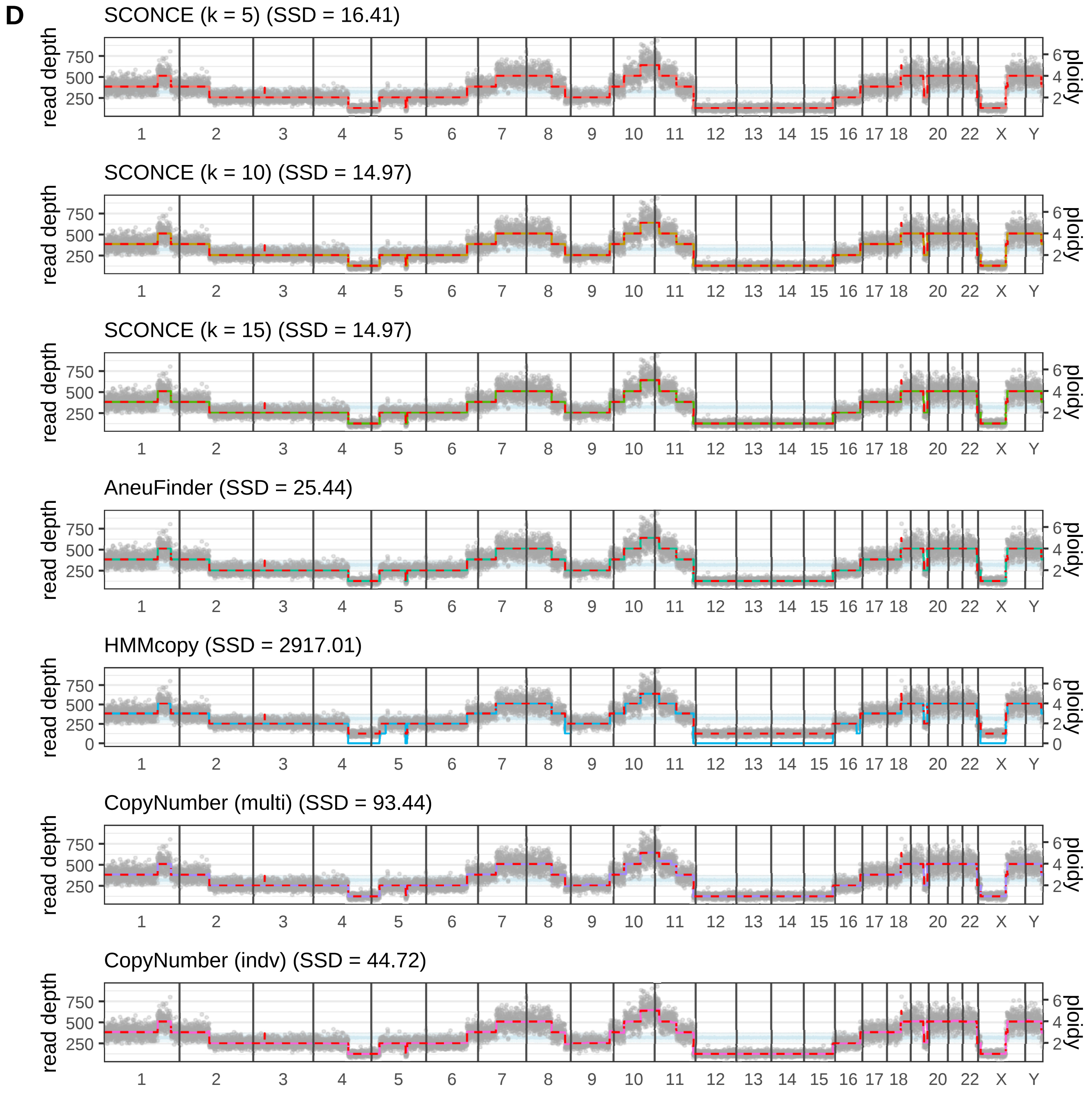


Fig. S3D: Genome wide decoding from cell 40 in Simulation Set D (mainly insertions under the line segment model).

Figure S3: As in Figure 3, genomic window is plotted along the x-axis, per window read depth is shown along the left y-axis, and ploidy is plotted along the right y-axis. Black vertical lines denote chromosome boundaries, gray dots represent observed tumor read depth in each window, the red dotted line denotes the true ploidy from simulation, the light blue line shows the mean diploid read count, the light blue band shows $\pm 1$ standard deviation in the diploid read count, and the colored lines denote the copy number decoding from each method. In each simulation condition, SCONCE consistently decodes the correct copy number state. Of note, as in panels A and B, the value of $k$ must be set high enough to allow the HMM to properly decode the true copy number state. Additionally, SCONCE repeatedly correctly identifies regions with no read coverage, as opposed to HMMcopy and CopyNumber (panels A and C). Futhermore, in panel C, AneuFinder incorrectly doubles all ploidy estimates, while SCONCE does not.

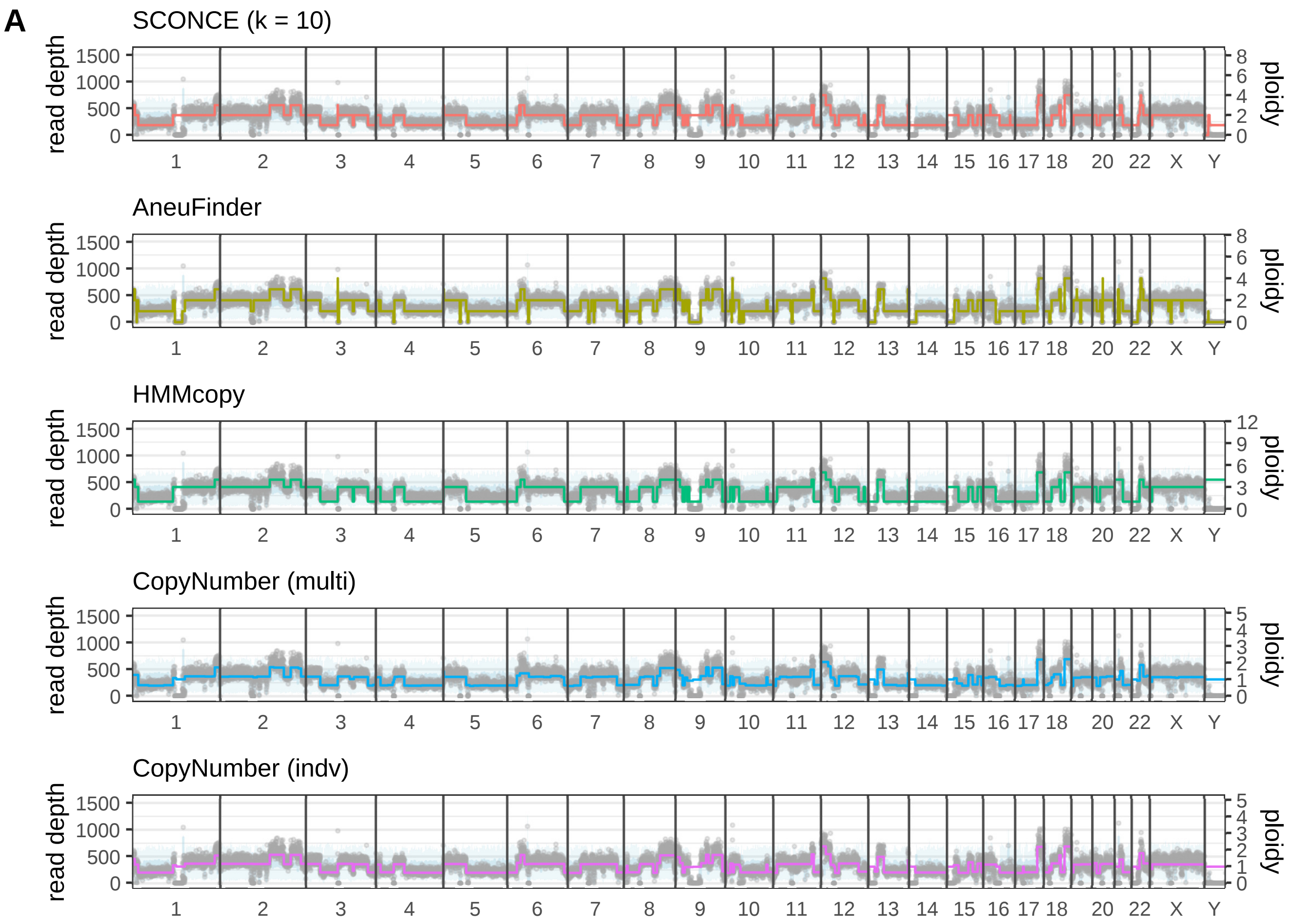


Fig. S4A: Genome wide decoding from cell SRR054570.

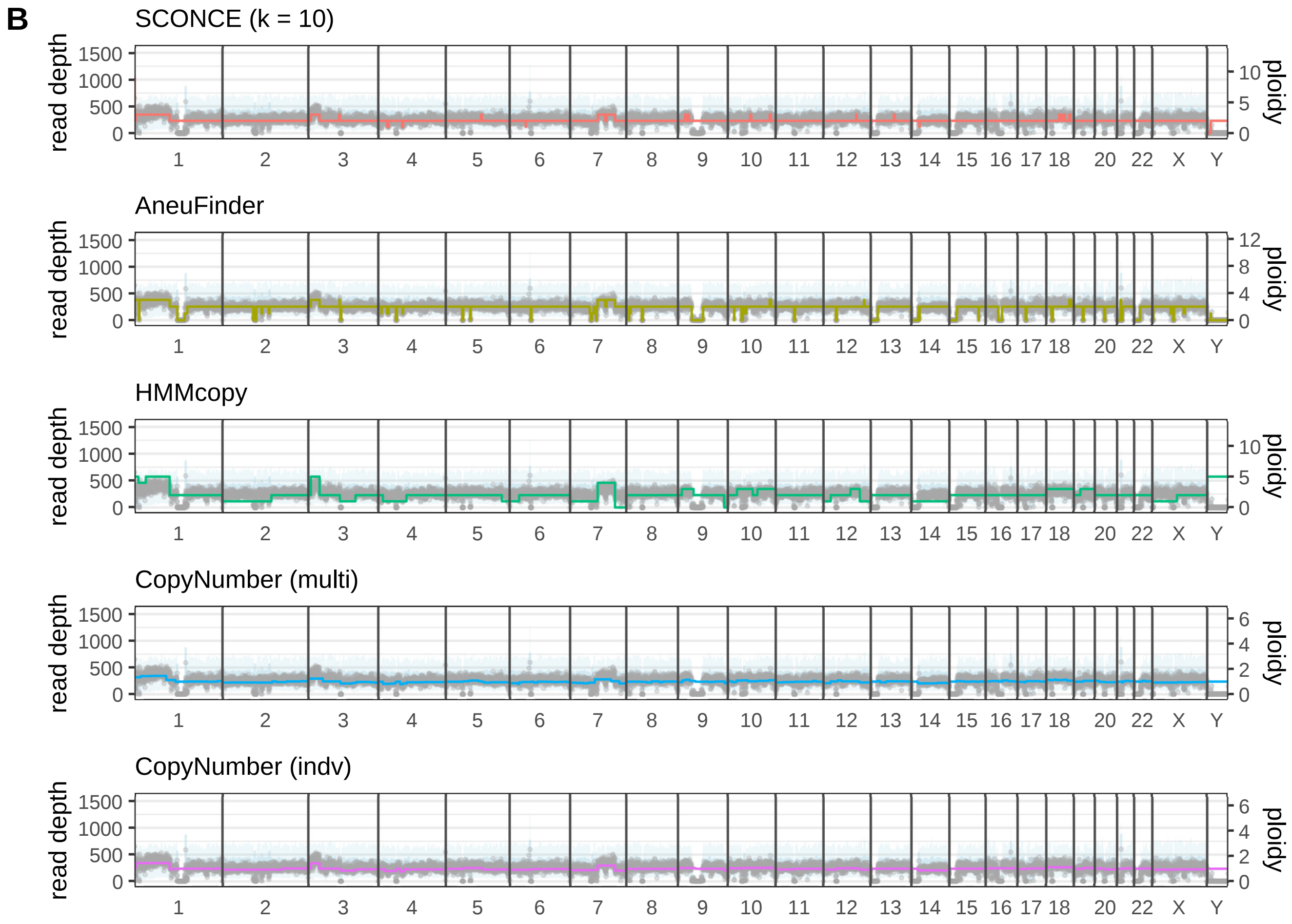


Fig. S4B: Genome wide decoding from cell SRR053675.

Figure S4: As in Figure 3 and Supplemental Figure S3, genomic window is plotted along the x-axis, read depth along the left y-axis, and ploidy along the right y-axis, with black vertical lines denoting chromosome boundaries. For each window, gray dots show observed tumor read depth, the light blue line shows the mean diploid read count, and the light blue band shows $\pm 1$ standard deviation in the diploid read count. Colored lines denote the copy number decoding from each method. Recapitulating the simulation results, SCONCE is more sensitive to small CNAs than AneuFinder (chromosomes 9, 10, 12, 13, and 18 in panel B). Additionally, SCONCE uses the null diploid model to predict no changes in copy number in hard to sequence and map regions that have no observed diploid or tumor reads (chromosomes 1, 9, 13-16, 21, and 22 in panel A).
